# Contrasting beta diversity and functional composition of aquatic insect communities across local to regional scales in Amazonian streams

**DOI:** 10.1101/2020.09.14.297077

**Authors:** Gilberto Nicacio, Erlane José Cunha, Neusa Hamada, Leandro Juen

## Abstract

We investigated how components of beta diversity (i.e., the turnover and nestedness and functional compositional) aquatic insect assemblages change among sites and are influenced by environmental and spatial drivers. For this, we analyzed beta-diversity and functional composition of Ephemeroptera, Plecoptera, and Trichoptera in 16 streams in two Amazonian basins with distinct environmental conditions (the Carajás and Tapajós regions). We performed Multiple regression on dissimilarity matrices (MRM) and Procrustes analysis to test spatial and environmental influences on the taxonomic and functional composition of communities. Community dissimilarity was most related to variations in geographic distance and topography, which highlighted the environmental distances shaping the communities. Variation in functional composition could be mostly attributed to the replacement of species by those with similar traits, indicating trait convergence among communities. Environmental predictors best-explained species replacement and trait congruence within and between the regions evaluated. In summary, among communities with different taxonomic compositions, the high species replacement observed appears to be leading them to have similar community structure, with species having the same functional composition, even in communities separated by both small and large geographic distances.

## 1. Introduction

Habitat structure and spatial distribution have long been considered the main determinants of species diversity and distribution (Tilman, 2004). Models of how niche-related mechanisms are affected by habitat conditions also have been related to species interactions and ecosystem functioning (Dangles & Malmqvist, 2004). On the other hand, neutral models of community ecology have assumed a stochastic perspective for dispersal-related processes, speciation, and extinction of organisms, in which all the species in an assemblage consist of ecologically equivalent individuals from the regional species pool (Stephen P Hubbell, 2001; Willig et al., 2003). Then, the effects of geographic distance and dispersal limitations have supported many neutral models of community structuring (Astorga et al., 2012).

To date, both types of determinants have been used in many studies to explain the mechanisms shaping community composition with regard to the geographic distances among communities and the specific dispersal limitations of the species considered (Moritz et al., 2013; Willig et al., 2003). Together, these theories have contributed to many models that have been applied to describe patterns in species distribution and community composition along environmental and spatial gradients (Blanchet et al., 2008). Although the primary assumption used in traditional niche-based approaches ignores neutral effects and emphasizes deterministic processes, many studies have also considered the relative importance of stochasticity in generating changes in community composition (beta-diversity) along environmental gradients (Chase & Myers, 2011).

Recently, to understand differences among biological communities, the variation in species composition among communities has been studied by examining the responses of beta-diversity by partitioning it into its replacement and richness difference components to find patterns along environmental and spatial gradients (Baselga, 2010). The response to these gradients assumes that community composition can be related to the difference between the number of species (i.e., richness difference), whereas contributions of turnover (i.e., species replacement), which refers to the addition of new species to the communities (Carvalho et al., 2012).

When differences in richness between communities are the main component of beta-diversity, the loss or gain of species should also result in changes in ecosystem function resulting from the homogenization of trait diversity and composition at low levels of taxonomic diversity (STEPHEN P. Hubbell, 2005). In contrast, turnover among highly diverse assemblages should buffer communities due to them having greater levels of complementarity and redundancy, and therefore these communities should be less strongly affected by changes in trait richness resulting from the loss or gain of a single species (STEPHEN P. Hubbell, 2005; Schriever et al., 2015). Then, assessing these patterns may help to disentangle the ecological processes structuring the taxonomic and functional composition of communities in response to deterministic and stochastic processes (Stegen & Hurlbert, 2011).

Contrasting taxonomic and functional beta-diversity can disentangle levels of spatial scales that determine the relative contribution of environmental or dispersal-based processes for community assembly (Sokol et al., 2011). For instance, in lotic ecosystems, it is expected that environmental gradients at local (stream) and regional (large basins) scales generate distinct patterns in taxonomic and functional beta-diversity within and among regions (Nekola & White, 1999; Soininen et al., 2007). Hence, if communities have high levels of functional beta-diversity due to the high frequency of replacement of species with unique traits, this is indicative of niche differentiation among functionally different communities (Villéger et al., 2013). In contrast, low functional beta-diversity may indicate that communities are functionally similar in the trait compositions of their species. If communities have high functional beta-diversity in their nested components, then different niche-filtering mechanisms may be shaping their functional compositions (Villéger et al., 2013).

Considering riverine communities increasing the spatial scale (e.g., from basin-scale to large interfluves), high extinction-colonization dynamics should play the main role in determining richness differences and are expected to promote nestedness between communities based on their species’ traits and dispersal abilities at large spatial scales (Miyazono & Taylor, 2013). In contrast, low extinction-colonization dynamics should play the main role in determining the beta-diversity (as the turnover component) within the basin-scale and are expected as a result of local dispersal rate (López-Delgado et al., 2020).

Therefore, here we investigate how components of beta diversity (i.e., the turnover and nestedness and functional compositional) aquatic insect assemblages change among sites and are influenced by environmental and spatial drivers. For this, we analyzed the beta-diversity patterns among aquatic insect communities (Ephemeroptera, Plecoptera, and Trichoptera) from forested streams in two distinct environmental contexts (regions) and evaluated species replacement and richness differences along communities and geographical scales. In order to examine beta-diversity patterns and functional composition of aquatic insect communities in response to environmental predictors at different spatial resolutions, this study addressed whether (i) that beta-diversity patterns at local scales (within regions) is due to species replacement occurring in response to the environment or dispersal processes; (ii) At larger geographic scales (e.g., between Amazonian interfluves), dispersal processes should play the main role in determining richness differences and were thus expected to promote nestedness between communities based on their species’ traits and dispersal abilities at large spatial scales.

## 2. Materials and methods

### 2.1. Study area

We evaluated 16 sites (first- and second-order streams) located in two pristine forested regions in the Floresta Nacional do Tapajós and Floresta Nacional de Carajás, both in Pará State, Brazil (Figure 1). Sampling was done in June 2015 and September/October 2015. The sampled sites at Flona Tapajós were streams with sandy beds distributed in lowland riverine networks. In contrast, the sites in the Flona Carajás region were streams with rocky beds, where the landscape has a high elevation range, from sea level at sites adjacent to the Amazon River to 600m above sea level in the Serra dos Carajás uplands.

**Figure 1.**
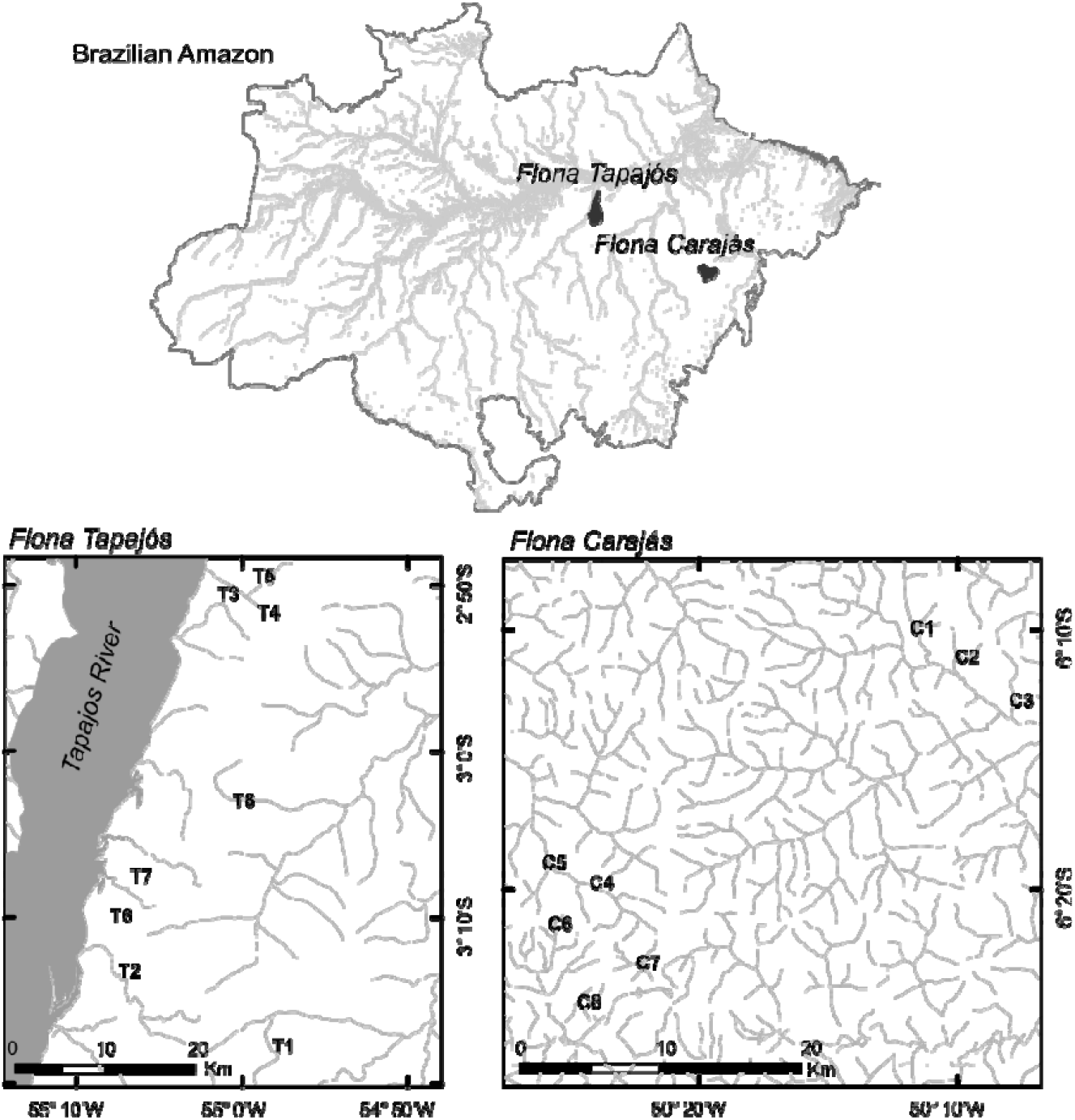
Study area: Floresta Nacional do Tapajós (Flona Tapajós) at Santarém/Belterra and Floresta Nacional de Carajás (Flona Tapajós) at Parauapebas/Canaã dos Carajás, Pará State, Brazil.

### 2.2 Field sampling and sample processing

Ephemeroptera, Plecoptera, and Trichoptera (EPT) were collected using a circular net (diameter = 18cm; mesh size = 250µm). At each stream, 20 benthic subsamples were collected systematically from all available instream’s habitats (e.g., cobble, woody debris, vegetated banks, submerged leaves, sand, and other fine sediments) by kicking the substrate into the circular dip nets. We performed the same sampling effort for all sites in each stream from the downstream end of the reach to 150 m upstream. Specimens were sorted in the field and preserved in 85% ethanol. Specimens were identified to the genus level when possible using the available literature, but while acknowledging the limited knowledge available on the Neotropical fauna (Domínguez et al. 2006; Hamada et al. 2014). The specimens were stored in the Zoological Collection at the Universidade Federal do Pará, Belém, Brazil.

### 2.3 Environmental data

We measured physical stream characteristics (mean wetted width/depth, mean substrate diameter, and elevation) from each stream along a 150-m-long stretch subdivided into 10 continuous sections, each 15 m long and with 11 cross-sectional transects taken in each stream (Kaufmann et al., 1999). In addition, variables from water at each stream cross-sectional transects from downstream, midstream, and upstream. These included dissolved oxygen content, conductivity, pH, and temperature (Table 1, Table S1 in Electronic Supplementary Material).

**Table 1.** Summary of environmental variables considered as predictors for variation in community composition of aquatic insects from streams at Floresta Nacional do Tapajós and Floresta Nacional de Carajás, Pará, Brazil.

| Name | Code | Average | SD | min | max |
| --- | --- | --- | --- | --- | --- |
| Mean substrate diameter (mm) | xsub | 44.248 | 11.962 | 20.000 | 61.905 |
| Mean wetted width / depth (m/m) | xwd | 9.501 | 2.862 | 5.381 | 14.906 |
| pH | pH | 5.897 | 1.190 | 4.493 | 8.120 |
| Electrical conductivity ($\mu\text{S}/\text{cm}$) | Cond | 22.408 | 18.063 | 3.000 | 78.000 |
| Temperature ($^{\circ}\text{C}$) | T | 23.977 | 1.636 | 20.900 | 25.833 |
| Dissolved oxygen (mg/L) | OD | 6.854 | 2.861 | 4.133 | 16.467 |

### 2.4 Functional trait composition of Ephemeroptera, Plecoptera, and Trichoptera (EPT)

To investigate the patterns in functional composition, a categorical genus-traits matrix (of EPT functional composition) was developed using traits from the available literature (see References in Electronic Supplementary Material). Six trait groups were then computed to express 20 trait states for each taxon identified using data available in the current literature. Then, trophic traits (i.e., ‘food’ and ‘guilds’), respiration modes, morphological adaptations (body shape and specific adaptations), and mobility were grouped (see the trait matrices in Tables S2 and S3 in Electronic Supplementary Material).

### 2.5 Statistical analyses

We performed principal component analysis (PCA) on the environmental variables using the correlation matrix to summarize environmental patterns within and between regions. Then, to represent the influence of spatial patterns on differences in the sampled communities, we used the pairwise Euclidean distances between sites calculated from their geographic coordinates (hereinafter referred to geographic distance).

To investigate our hypotheses, taxonomic and functional beta-diversity values were estimated from dissimilarity matrices based on presence/absence methods, and their components were computed following the approaches proposed by Baselga (2010) and Villéger et al. (2013). Using Baselga’s (2010) approach, we calculated the Sorensen-based multiple-site dissimilarity (βsor), the Simpson-based multiple-site dissimilarity (βsim), and the nestedness-based multiple-site dissimilarity (βnes).

Distance matrices were produced for analyses both within and between regions based on their overall variation in community composition (beta-diversity: βsor), species replacement (turnover: βsim), and richness difference (nestedness: βnes). Then, for the decomposition of functional beta-diversity (i.e., the dissimilarity in functional composition among communities), a functional turnover and functional nestedness component were calculated as a functional distance matrix between each pair of species using Gower’s distance. In addition, non-metric multidimensional scaling (nMDS) was performed to summarise patterns in community composition.

To test the hypothesis that geographic distance affects community dissimilarity, the community dataset was evaluated as two groups (the Carajás and Tapajós regions). Then, Multiple regression on dissimilarity matrices (MRM) was then performed using the Sorensen index-based distances to test the relationships among beta-diversity and the spatial and environmental predictors (based on the Euclidean distance matrix).

To investigate the patterns in functional composition among communities, we performed the Hill and Smith ordination method on the categorical trait matrix weighted based on scores from the abundance matrix to obtain trait scores that represented spatial and functional variations (Dray et al., 2014). Procrustes analysis was then performed to estimate the degree of association between traits in communities from the Carajás and Tapajós regions. This analysis is used to assess similarities in trait compositions that are weighted by the distinct abundances of the assemblages in each stream site. Procrustes analysis aims to find matches between ordinations and associated matrices and produces an m^2^-statistic that is transformed into an r-statistic (r = square root of (1-m^2^)). The r-statistic indicates the strength of the correlation between the results of two ordinations (Peres-Neto & Jackson, 2001).

We then performed MRM with Gower’s dissimilarity index values to test the effects of the environmental and spatial filters on the functional composition of the communities, as measured by the community-level weighted means of trait values (Lavorel et al., 2007). We performed all statistical analyses with functions from packages vegan, ade4, ecodist, and labdsv in R version 3.3.0 (R Core Team 2016).

## 3. Results

### 3.1 Summary of environmental variables

PCA results showed that the streams in both regions (Flona Carajás and Tapajós) formed heterogeneous spatial groups from one another, differentiated by the values of the environmental variables in each. Flona Carajás is a region with mostly bedrock-bottomed streams that typically had high pH values, representing habitat with alkaline waters, as well as low temperatures and high electrical conductivity values. In contrast, habitats in Flona Tapajós were streams with sandy bed that had more acidic waters and a lower substrate size (Figure 2).

**Figure 2.**
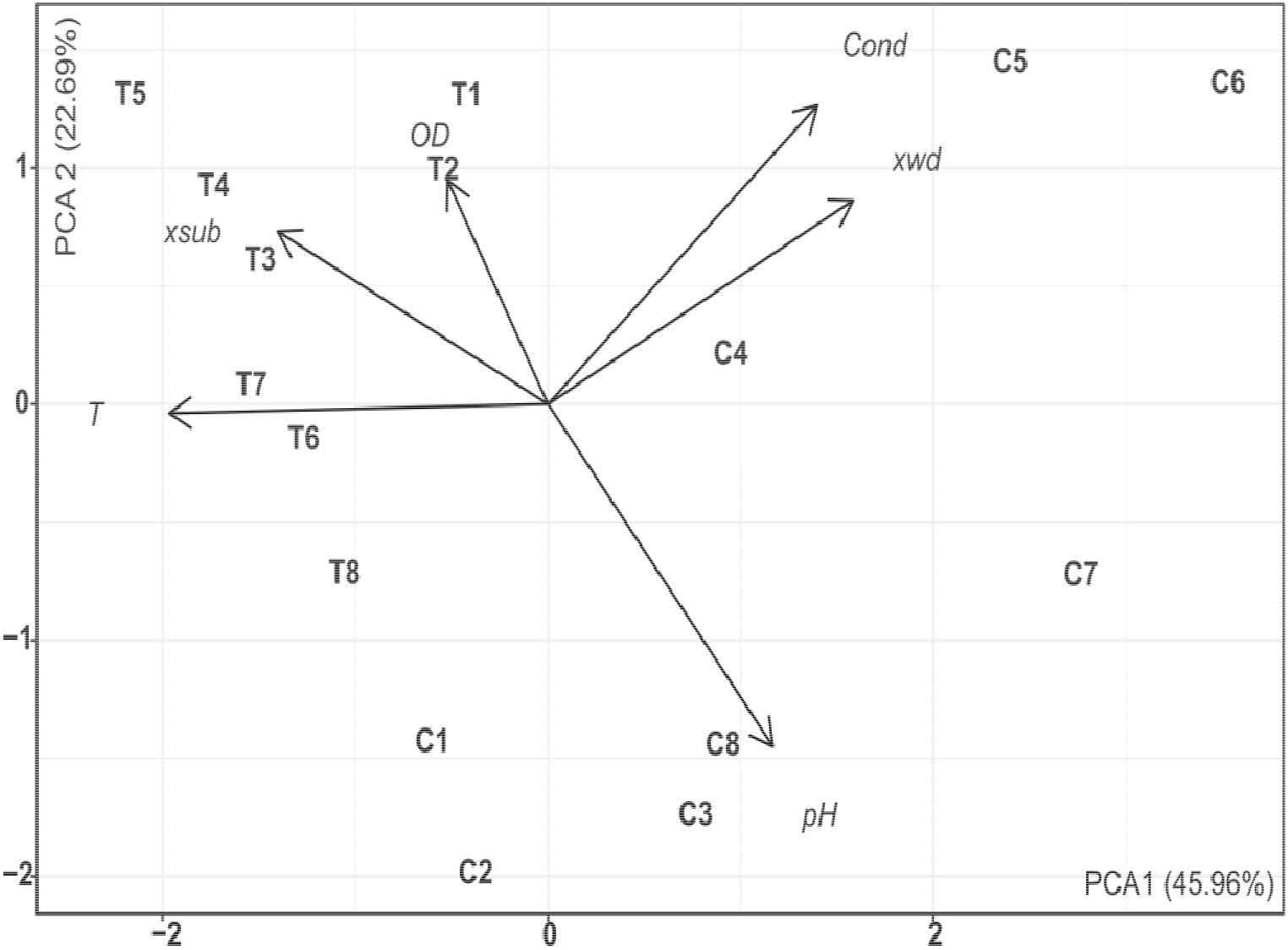
Plot of principal component analysis (PCA): Environmental variables from streams at Floresta Nacional de Carajás (C1-C8) and Floresta Nacional do Tapajós (T1-T8), Pará State, Brazil. Variable codes: mean substrate diameter (xsub), mean wetted width/depth (xwd), negative log hydrogen ion concentration (pH), electrical conductivity (Cond), temperature (T), and dissolved oxygen (OD).

### 3.2 Overall community composition

We recorded 6,704 immature specimens of Ephemeroptera, Plecoptera, and Trichoptera (EPT); 3,717 specimens were collected in Flona Carajás and 2,987 were collected in Flona Tapajós (Table S4 in Electronic Supplementary Material). We identified 53 genera, which represented 30 Ephemeroptera, 2 Plecoptera, and 21 Trichoptera genera. The overall genus richness in the sampled regions averaged 19 genera per stream in Flona Carajás and 24 genera per stream in Flona Tapajós.

Each studied region had its unique EPT genera, and the difference in richness between regions was due to 17 unique genera in Flona Carajás and 14 unique genera in Flona Tapajós. The following genera were exclusively collected in Flona Carajás: *Atopsyche, Brasilocaenis, Caenis, Callibaetis, Hydrosmilodon, Leentvaaria, Leptohyphes, Leptohyphodes, Notalina, Notidobiella, Paracloeodes, Paramaka, Polycentropus, Terpides, Traverhyphes, Tricorythodes*, and *Ulmeritoides*. In contrast, the following genera were exclusively collected in Flona Tapajós: *Amazonatolica, Americabaetis, Apobaetis, Aturbina, Austrotinodes, Campsurus, Cloeodes, Cryptonympha, Cyrnellus, Leptoceridae sp*., *Hydrosmilodon, Simothraulopsis, Tricorythopsis*, and *Waltzoyphius*.

### 3.3 Components of taxonomic and functional beta-diversity

Overall, when assessing multiple-site dissimilarities among streams, the complete dataset showed that there was high variation in community composition (βsor = 0.751), and when we quantified the contributing causes, we found that species replacement had a strong effect (βsim = 0.677) while richness differences had only a weak effect (βnes = 0.073). The estimated overall beta-diversity within each region, Carajás (βsor = 0.594) and Tapajós (βsor = 0.502), indicated that beta-diversity contributed similarly to community dissimilarity within both areas.

When we partitioned this overall beta-diversity into its components, species replacement (i.e., turnover) and richness difference, turnover was the process that was responsible for most of the beta-diversity in streams in both the Carajás (βsim = 0.506, βnes = 0.087) and Tapajós (βsim = 0.420, βnes = 0.083) regions. Considering only the dissimilarities among streams within each region, we detected weak responses of the beta-diversity components to geographic distance using pairwise measures (the Euclidean distances among streams).

For functional composition, we found quite a low function diversity overall (β_F_ = 0.417), and when we quantified the contributing causes of this, we found that species replacement had the strongest effect (Turnover _F_ = 0.300) and that richness differences had only a weak effect (Nestedness _F_ = 0.116).

### 3.3. Community responses to environmental and spatial variations

The non-metric multidimensional scaling (nMDS), based on the Sorensen index values for presence/absence data, highlighted the high variation in community composition in the studied systems, and showed that they could be separated into distinct spatial groups based on their regional distribution (Figure 3). Considering the species distributions within each region, we found greater dissimilarity in community composition among the streams in Flona Carajás, while the assemblage dissimilarities among streams in Flona Tapajós were lower.

**Figure 3.**
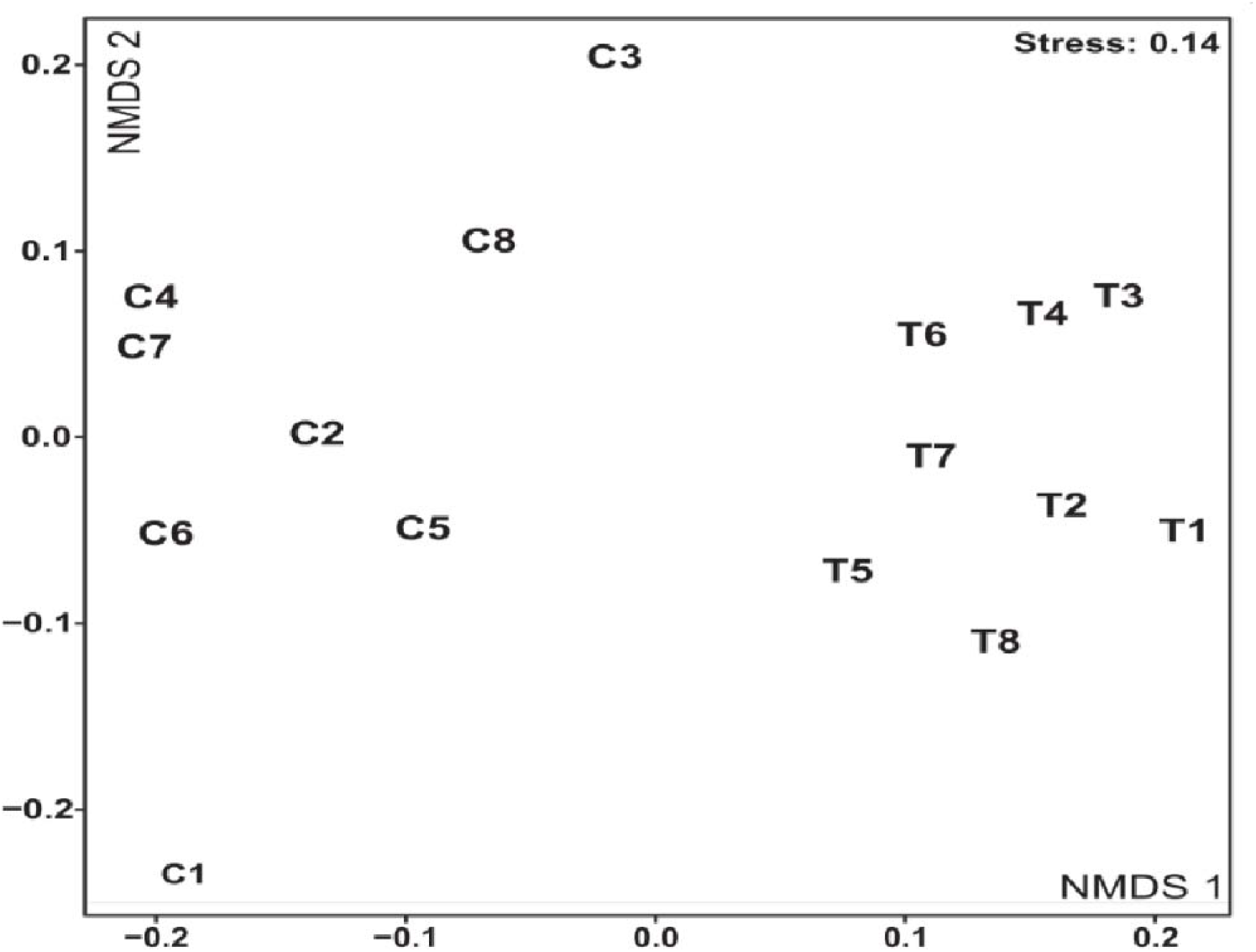
Plot of Non-metric multidimensional scaling (NMDS) - Sørensen dissimilarity: Ephemeroptera, Trichoptera, and Plecoptera assemblages from streams at Floresta Nacional de Carajás (C1-C8) and Floresta Nacional do Tapajós (T1-T8), Pará State, Brazil

The overall functional composition was remarkably similar between the two regions, as the congruence found in the functional matrices. Procrustes analysis, based on scores calculated by the Hill and Smith method (functional traits weighted by community abundance), resulted in a significant congruence being found between the functional composition of stream communities in Flona Carajás and Tapajós (m^2^ = 0.009, r = 0.949, p < 0.001).

Results of multiple regression on dissimilarity matrices (MRM) demonstrated the individual relationships among the environmental variables and the community matrix (Sorensen dissimilarity). The overall species replacement among streams was individually correlated with elevation, temperature, and pH. These variables were significantly correlated in the model that included geographic distance as an explanatory variable (Table 2). In contrast, we did not find any significant relationships between functional composition and environmental variables (R^2^ = 0.014, F = 1.740, p = 0.436) or geographic distance (R^2^ = 0.001, F = 1.128, p = 0.454) based on analyses of dissimilarity matrices (Gower distances).

**Table 2.** Results for multiple regression on distance matrices. Relationships between the taxonomic β‐diversity (Sørensen distance) and the explanatory variables for aquatic insect assemblages (Ephemeroptera, Trichoptera, Plecoptera) from streams at Floresta Nacional do Tapajós and Floresta Nacional de Carajás, Pará, Brazil.

| Variables | Code | R2 | F-test | p |
| --- | --- | --- | --- | --- |
| *Individual response* |  |  |  |  |
| Geographic distance | xy | 0.497 | 116.774 | 0.001 |
| Mean wetted width / depth (m/m) | xwd | 0.001 | 0.072 | 0.812 |
| Mean substrate diameter (mm) | xsub | 0.001 | 0.050 | 0.842 |
| Elevation | elev | 0.438 | 92.160 | 0.001 |
| Electrical conductivity ($\mu\text{S}/\text{cm}$) | cond | 0.027 | 3.240 | 0.129 |
| Dissolved oxygen (mg/L) | OD | 0.002 | 0.264 | 0.680 |
| Temperature ($^{\circ}\text{C}$) | T | 0.112 | 14.846 | 0.006 |
| pH | pH | 0.381 | 72.522 | 0.001 |
| *Model with environmental and spatial distances* |  |  |  |  |
| xy + elev + T + pH | - | 0.511 | 60.865 | 0.001 |

## 4. Discussion

We found support for our first hypothesis that beta-diversity patterns (i.e., differences in species composition) at the scale of local habitats (i.e., within regions) were mostly related to species replacement. The richness differences expected by our second hypothesis contributed the least to patterns in beta-diversity, which indicated that the expected dispersal dynamics had only small effects at larger geographic scales. The high frequency of species replacement observed indicated that species distributions were mainly driven by geographic distances and only weakly impacted by environmental influences, as was expected based on species sorting (i.e., environmental filtering) that emerge in response to dispersal processes along environmental gradients. Then, we found support the congruence of functional traits among streams (low functional beta-diversity by species replacement) also supported the hypothesis that neutral dynamics are important in the emergence of functional equivalence. This could occur because, although the high species turnover may be leading to dissimilarities in species composition among the studied communities, species are also being replaced by other species that perform the same functions in different streams.

Examining species replacement allowed us to identify the key environmental processes involved in determining community dissimilarity. Specific regional contexts may be controlling the distributions of aquatic insects along the environmental gradients within and between the Flona Carajás and Tapajós regions. Considering the regional specificity of the assemblages found, the most important explanatory variables determining species replacement between the two regions were geographic distance and elevation. The pH can be highlighted as the local environmental predictor for beta-diversity, which commonly represent good predictor for the distributions of Ephemeroptera, Plecoptera, and Trichoptera (Bispo et al., 2006). In addition to the spatial distribution of streams between the regions, species replacement was most strongly structured by the regional conditions that were presented in the PCA results.

The observed patterns in streams from Flona Carajás can be a combination of both effects of high-altitude catchments (i.e. spatial influences) and local environmental context characterized as having high values of electrical conductivity and slightly neutral water. In contrast, the habitats in streams in Flona Tapajós had remarkably low electrical conductivity values and acidic water and were distributed in lowland drainages. Site-specific attributes like these are known to create habitat heterogeneity and contribute significantly to species turnover within regional contexts (Datry et al., 2016). Highlighting the influence of elevation, the high values of overall beta-diversity and species replacement observed for the Carajás region can be explained by the weak influence of dispersal limitation on the structuring of these assemblages. Because high elevations can act as barriers to dispersal, they can lead to selective differentiation among the fauna in different streams, as was suggested by the high turnover patterns found, especially in the higher-elevation Carajás region.

Specific environmental gradients are known to control community composition and determine the similarities among assemblages of aquatic fauna (Heino et al., 2003; López-Delgado et al., 2020). Unexpectedly, we did not find relationships between local stream conditions (e.g., substrate size, electrical conductivity, and stream width) and community dissimilarity, which could be related to different factors acting more strongly in structuring local patterns than those acting in regional species replacement (Hamerlík et al., 2014). Variations in community composition caused by species replacement patterns were expected in the studied communities since they were in different environmental contexts in different regions, as was expressed in the results of the regression analysis methods applied. Unexpectedly, congruent in the functional compositions of streams was found between the two regions. This result suggests, that at the regional scale, differences in environmental conditions may be more important to affect taxonomic composition, and thus the functional composition may respond through the convergence of traits between regions. which is related to the functional redundancy at the scale of the regional species pool (Finn & Poff, 2005).

When partitioning the taxonomic composition and considering the spatial patterns among assemblages using ordination plots, some patterns became evident. For instance, the nMDS results explained the distributions of genera between the regions, which were associated with the environmental contributions to differences in community composition (Costa & Melo, 2008; Patrick & Swan, 2011). When describing assemblage distributions, we found that each insect order made a distinct contribution to species replacement. As a result, we found that the distribution and species replacement of Ephemeroptera assemblages (e.g., *Callibaetis, Brasilocaenis, Hydrosmilodon, and Ulmeritoides*) were the most important factors leading to differences in assemblages among streams in the Carajás region. In contrast, beta-diversity patterns for most Trichoptera genera were more evident in habitats in the Tapajós region. These observations highlight typical patterns in stream environments along black-water drainages, where specific habitats are created for different aquatic biota.

The functional composition in Tapajós region was mostly comprised of collector-gatherers, which were present in high relative proportions of the communities (e.g., *Miroculis*); this suggests that these organisms are associated with the processing of coarse (CPOM) and fine particulate organic matter (FPOM), which, in turn, may be used as food by these insects (Baptista et al., 2006). Furthermore, shredders (e.g., *Nectopsyche* and *Phylloicus*) contributed the most to the functional patterns observed among most streams in Tapajós. The Tapajós region is known to receive high inputs of litter material produced in riparian forests, which leads to large amounts of leaf debris being present that decomposes and creates unique conditions for aquatic fauna in these streams (Bonetto & Sioli, 1975).

In our results, the absence of intermediate regions only allowed us to explore regional patterns in beta-diversity components and functional composition. Therefore, we must acknowledge the spatial limitations of our study and our inability to state broad generalizations for application to larger scales than those examined herein. Nevertheless, the beta-diversity patterns and the high functional congruence found between sites can be credited to the framework applied in this study, due to the relatively small number of streams assessed and the uneven distances they had from each other and between each region (Heino, 2009b). This suggests that adding more sites along more pronounced and broader environmental and spatial gradients would increase the variability in the species functional organization and community dissimilarity that would be detected, since large-scale variation increases the heterogeneity of stream habitats (Heino, 2009a; Ricotta & Burrascano, 2008).

Despite the above limitations, we found little evidence of environmental influences on richness differences (nestedness) and strong evidence of environmental influences on species replacement (spatial turnover). Therefore, some ecological implications from our findings should be highlighted that showed the contributions of environmental deterministic processes and the absence of dispersal limitation effects in some contexts. For instance, high variation in environmental conditions within and between regions was demonstrated in the PCA, and this pattern was supported by the community dissimilarity observed when there were low richness differences but strong turnover effects among streams, reflecting the habitat heterogeneity across the landscape between regions.

## 5. Conclusion

Spatial and environmental variation represented key factors for assemblage composition of Ephemeroptera, Plecoptera, and Trichoptera. The observed patterns can be a combination of high replacement and low species loss species associated to congruence in functional composition. Moreover, species replacement were the most strongly related to variations in landscape features (geographic and environmental); however, the functional composition did not diverge concurrently with taxonomic dissimilarity between the regions, which may be related to there being convergent regional trait pools due to similarities in community dynamics and microhabitat distributions between regions. Thus, when replacement of functionally similar species is high, the regional and local environmental conditions may be able to maintain functional congruence in similar habitats, even those with different taxonomic compositions.

