## Supplementary material for "Contrasting beta diversity and functional composition of aquatic insect communities across local to regional scales in Amazonian streams": Table S1

**Table S1.** Summary of environmental variables measured from streams at Floresta Nacional do Tapajós and Floresta Nacional de Carajás, Pará State, Brazil.

| **Sites** | **Cond** | **OD** | **T** | **pH** | **xwd** | **xsub** | **Elevation** | **Longitude** | **Latitude** | **Region** |
| --- | --- | --- | --- | --- | --- | --- | --- | --- | --- | --- |
| C1 | 5.16 | 5.72 | 24.40 | 7.31 | 8.76 | 54.29 | 355 | -50.19310 | -6.16560 | Carajás |
| C2 | 3.00 | 6.34 | 24.54 | 7.20 | 7.25 | 35.24 | 399 | -50.16410 | -6.18399 | Carajás |
| C3 | 15.00 | 4.70 | 24.20 | 7.21 | 9.51 | 34.00 | 369 | -50.12920 | -6.19532 | Carajás |
| C4 | 30.00 | 6.04 | 22.50 | 6.63 | 10.04 | 52.38 | 489 | -50.41100 | -6.32901 | Carajás |
| C5 | 45.00 | 5.45 | 21.20 | 6.10 | 13.70 | 53.33 | 535 | -50.43790 | -6.33369 | Carajás |
| C6 | 48.00 | 6.54 | 20.90 | 6.38 | 14.91 | 32.38 | 559 | -50.42640 | -6.35537 | Carajás |
| C7 | 32.00 | 5.30 | 21.70 | 6.69 | 11.00 | 20.00 | 558 | -50.36950 | -6.38066 | Carajás |
| C8 | 10.00 | 6.98 | 23.00 | 8.12 | 10.16 | 40.00 | 275 | -50.40330 | -6.41573 | Carajás |
| T1 | 15.30 | 8.07 | 24.73 | 4.49 | 12.35 | 41.25 | 137 | -54.96360 | -3.29705 | Tapajós |
| T2 | 14.27 | 6.59 | 25.30 | 4.63 | 13.14 | 47.62 | 41 | -55.12620 | -3.20269 | Tapajós |
| T3 | 19.70 | 7.70 | 25.43 | 4.89 | 6.82 | 47.00 | 39 | -55.01300 | -2.83363 | Tapajós |
| T4 | 20.17 | 7.40 | 25.83 | 4.83 | 7.86 | 55.24 | 58 | -54.99530 | -2.84336 | Tapajós |
| T5 | 19.63 | 8.53 | 25.63 | 4.90 | 7.12 | 61.91 | 57 | -55.00190 | -2.83718 | Tapajós |
| T6 | 18.53 | 4.13 | 25.23 | 4.74 | 7.77 | 56.67 | 37 | -55.11670 | -3.14579 | Tapajós |
| T7 | 20.63 | 4.63 | 25.33 | 4.82 | 6.26 | 57.14 | 54 | -55.10650 | -3.13715 | Tapajós |
| T8 | 12.13 | 5.67 | 23.70 | 5.40 | 5.38 | 49.52 | 129 | -55.00290 | -3.04941 | Tapajós |

Variable codes: Mean substrate diameter (xsub), mean wetted width/depth (xwd), negative log hydrogen ion concentration (pH), electrical conductivity (Cond), temperature (T), dissolved oxygen (OD).

**Table S2.** Traits and their modalities describing functional traits of aquatic insect taxa computed to aquatic assemblages from streams at Floresta Nacional do Tapajós, Pará State, Brazil.

| **Traits** | **Modalities** | **Codes** | **N** |
| --- | --- | --- | --- |
| Food | Coarse particulate organic matter | CPOM | 10 |
|  | Fine particulate organic matter | FPOM | 36 |
|  | Macroinvertebrates | MaIn | 7 |
| Guild | Collectors/filters | CF | 8 |
|  | Collectors/gatherers | CG | 30 |
|  | Predator | PR | 7 |
|  | Shredder | SH | 8 |
| Respiration | Gill | gill | 37 |
|  | Tegument | teg | 16 |
| Body shape | Cylindrical | cyl | 22 |
|  | Flattened | flatt | 19 |
|  | Streamlined | stre | 12 |
| Specific adaptation to flow constraints | Anal hooks | AH | 10 |
|  | Mineral material (case) | MM | 2 |
|  | Silk gland | SG | 8 |
|  | Tarsal hooks | TH | 33 |
| Locomotion and relation to substrate | Crawler | CL | 25 |
|  | Full water swimmer | SwW | 9 |
|  | Temporally attached | TA | 19 |

* Trait groups and their modalities for each taxon were computed from available studies considering the limited knowledge on functional traits available for Neotropical fauna (e.g. Cummins et al., 2005; Tomanova et al., 2006; Tomanova & Usseglio-Polatera, 2007; Colzani et al., 2013; Castro et al., 2017). In addition, we compared our categorical matrix of species traits with prior studies that evaluated relationships between biological attributes of aquatic insects linked and the environment from temperate streams (Cummins, 1973; Finn & Poff, 2005; Poff et al., 2006; Merritt & Cummins, 2007; Merritt et al., 2008).

**Table S3.** Codes for trait matrix of Ephemeroptera, Plecoptera and Trichoptera assemblages from streams at Floresta Nacional de Carajás and Floresta Nacional do Tapajós, Pará State, Brazil.

| **Order** | **Family** | ***Genus*** | **Food** | **Guild** | **Respiration** | **Body shape** | **Specific adaptation**  **to flow constraints** | **Locomotion and**  **Relation to substrate** |
| --- | --- | --- | --- | --- | --- | --- | --- | --- |
| Ephemeroptera | Baetidae | *Americabaetis* | FPOM | CG | gill | stre | TH | SwW |
| Ephemeroptera | Baetidae | *Apobaetis* | FPOM | CG | gill | stre | TH | SwW |
| Ephemeroptera | Baetidae | *Aturbina* | FPOM | CG | gill | stre | TH | SwW |
| Ephemeroptera | Baetidae | *Callibaetis* | FPOM | CG | gill | stre | TH | CL |
| Ephemeroptera | Baetidae | *Callibaetoides* | FPOM | CG | gill | stre | TH | CL |
| Ephemeroptera | Baetidae | *Cloeodes* | FPOM | CG | gill | stre | TH | CL |
| Ephemeroptera | Baetidae | *Cryptonympha* | FPOM | CG | gill | stre | TH | CL |
| Ephemeroptera | Baetidae | *Paracloeodes* | FPOM | CG | gill | stre | TH | CL |
| Ephemeroptera | Baetidae | *Waltzoyphius* | FPOM | CG | gill | stre | TH | TA |
| Ephemeroptera | Baetidae | *Zelusia* | FPOM | CG | gill | stre | TH | TA |
| Ephemeroptera | Caenidae | *Brasilocaenis* | FPOM | CG | gill | stre | TH | CL |
| Ephemeroptera | Caenidae | *Caenis* | FPOM | CG | gill | stre | TH | CL |
| Ephemeroptera | Euthyplociidae | *Campylocia* | FPOM | CF | gill | flatt | TH | CL |
| Ephemeroptera | Leptohyphidae | *Amanahyphes* | FPOM | CG | gill | flatt | TH | SwW |
| Ephemeroptera | Leptohyphidae | *Leptohyphes* | FPOM | CG | gill | flatt | TH | TA |
| Ephemeroptera | Leptohyphidae | *Leptohyphodes* | FPOM | CG | gill | flatt | TH | TA |
| Ephemeroptera | Leptohyphidae | *Traverhyphes* | FPOM | CG | gill | flatt | TH | TA |
| Ephemeroptera | Leptohyphidae | *Tricorythodes* | FPOM | CG | gill | flatt | TH | TA |
| Ephemeroptera | Leptohyphidae | *Tricorythopsis* | FPOM | CG | gill | flatt | TH | TA |
| Ephemeroptera | Leptophlebiidae | *Askola* | FPOM | CG | gill | flatt | TH | SwW |
| Ephemeroptera | Leptophlebiidae | *Farrodes* | CPOM | CG | gill | flatt | TH | CL |
| Ephemeroptera | Leptophlebiidae | *Hagenulopsis* | FPOM | CG | gill | flatt | TH | CL |
| Ephemeroptera | Leptophlebiidae | *Hydrosmilodon* | FPOM | CG | gill | flatt | TH | TA |
| Ephemeroptera | Leptophlebiidae | *Leentvaaria* | FPOM | CG | gill | flatt | TH | TA |
| Ephemeroptera | Leptophlebiidae | *Miroculis* | CPOM | CG | gill | flatt | TH | TA |
| Ephemeroptera | Leptophlebiidae | *Paramaka* | FPOM | CG | gill | flatt | TH | TA |
| Ephemeroptera | Leptophlebiidae | *Simothraulopsis* | FPOM | CG | gill | flatt | TH | CL |
| Ephemeroptera | Leptophlebiidae | *Terpides* | CPOM | CG | gill | flatt | TH | TA |
| Ephemeroptera | Leptophlebiidae | *Ulmeritoides* | CPOM | CG | gill | flatt | TH | TA |
| Ephemeroptera | Polymitarcyidae | *Campsurus* | FPOM | CF | gill | cyl | TH | CL |
| Plecoptera | Perlidae | *Anacroneuria* | MaIn | PR | gill | flatt | TH | SwW |
| Plecoptera | Perlidae | *Macrogynoplax* | MaIn | PR | gill | flatt | TH | SwW |
| Trichoptera | Calamoceratidae | *Phylloicus* | CPOM | SH | gill | cyl | SG | CL |
| Trichoptera | Ecnomidae | *Austrotinodes* | FPOM | CF | teg | cyl | AH | SwW |
| Trichoptera | Helicopsychidae | *Helicopsyche* | FPOM | CG | teg | cyl | MM | CL |
| Trichoptera | Hydrobiosidae | *Atopsyche* | MaIn | PR | teg | cyl | TH | CL |
| Trichoptera | Hydropsychidae | *Leptonema* | FPOM | CF | gill | cyl | AH | TA |
| Trichoptera | Hydropsychidae | *Macronema* | FPOM | CF | gill | cyl | AH | TA |
| Trichoptera | Hydropsychidae | *Macrostemum* | FPOM | CF | gill | cyl | AH | TA |
| Trichoptera | Hydropsychidae | *Smicridea* | FPOM | CF | gill | cyl | AH | TA |
| Trichoptera | Leptoceridae | *Amazonatolica* | FPOM | SH | teg | cyl | SG | SwW |
| Trichoptera | Leptoceridae | Leptoceridae sp. | FPOM | SH | teg | cyl | SG | CL |
| Trichoptera | Leptoceridae | *Nectopsyche* | CPOM | SH | teg | cyl | SG | CL |
| Trichoptera | Leptoceridae | *Notalina* | CPOM | SH | teg | cyl | SG | CL |
| Trichoptera | Leptoceridae | *Oecetis* | CPOM | SH | teg | cyl | SG | CL |
| Trichoptera | Leptoceridae | *Triplectides* | CPOM | SH | teg | cyl | SG | TA |
| Trichoptera | Odontoceridae | *Marilia* | FPOM | CG | teg | cyl | MM | TA |
| Trichoptera | Philopotamidae | *Chimarra* | FPOM | CF | teg | cyl | AH | CL |
| Trichoptera | Polycentropodidae | *Cernotina* | MaIn | PR | teg | cyl | AH | CL |
| Trichoptera | Polycentropodidae | *Cyrnellus* | MaIn | PR | teg | cyl | AH | CL |
| Trichoptera | Polycentropodidae | *Polycentropus* | MaIn | PR | teg | cyl | AH | CL |
| Trichoptera | Polycentropodidae | *Polyplectropus* | MaIn | PR | teg | cyl | AH | CL |
| Trichoptera | Sericostomatidae | *Notidobiella* | CPOM | SH | teg | cyl | SG | CL |

**Table S4.** Abundance matrix of Ephemeroptera, Plecoptera and Trichoptera assemblages from streams at Floresta Nacional de Carajás (C1-C8) and Floresta Nacional do Tapajós (T1-T8), Pará State, Brazil.

| **Order** | **Family** | **Genus** | **C1** | **C2** | **C3** | **C4** | **C5** | **C6** | **C7** | **C8** | **T1** | **T2** | **T3** | **T4** | **T5** | **T6** | **T7** | **T8** |
| --- | --- | --- | --- | --- | --- | --- | --- | --- | --- | --- | --- | --- | --- | --- | --- | --- | --- | --- |
| Ephemeroptera | Baetidae | *Americabaetis* | 0 | 0 | 0 | 0 | 0 | 0 | 0 | 0 | 1 | 1 | 1 | 0 | 0 | 0 | 0 | 1 |
| Ephemeroptera | Baetidae | *Apobaetis* | 0 | 0 | 0 | 0 | 0 | 0 | 0 | 0 | 1 | 0 | 0 | 0 | 0 | 0 | 0 | 0 |
| Ephemeroptera | Baetidae | *Aturbina* | 0 | 0 | 0 | 0 | 0 | 0 | 0 | 0 | 0 | 1 | 0 | 0 | 0 | 0 | 0 | 1 |
| Ephemeroptera | Baetidae | *Callibaetis* | 0 | 0 | 0 | 0 | 0 | 2 | 1 | 0 | 0 | 0 | 0 | 0 | 0 | 0 | 0 | 1 |
| Ephemeroptera | Baetidae | *Callibaetoides* | 0 | 0 | 0 | 1 | 0 | 6 | 3 | 0 | 0 | 0 | 0 | 0 | 0 | 0 | 0 | 0 |
| Ephemeroptera | Baetidae | *Cloeodes* | 1 | 0 | 0 | 0 | 1 | 10 | 0 | 0 | 2 | 0 | 0 | 0 | 0 | 0 | 0 | 0 |
| Ephemeroptera | Baetidae | *Cryptonympha* | 0 | 0 | 0 | 0 | 0 | 0 | 0 | 0 | 0 | 0 | 2 | 0 | 0 | 0 | 0 | 0 |
| Ephemeroptera | Baetidae | *Paracloeodes* | 1 | 0 | 0 | 0 | 0 | 0 | 0 | 0 | 0 | 0 | 0 | 0 | 0 | 0 | 0 | 0 |
| Ephemeroptera | Baetidae | *Waltzoyphius* | 0 | 0 | 0 | 0 | 0 | 0 | 0 | 0 | 1 | 0 | 1 | 1 | 0 | 1 | 0 | 2 |
| Ephemeroptera | Baetidae | *Zelusia* | 4 | 6 | 1 | 4 | 7 | 6 | 9 | 3 | 4 | 1 | 14 | 5 | 2 | 5 | 1 | 15 |
| Ephemeroptera | Caenidae | *Brasilocaenis* | 6 | 0 | 0 | 0 | 1 | 0 | 0 | 0 | 0 | 0 | 0 | 0 | 0 | 0 | 0 | 0 |
| Ephemeroptera | Caenidae | *Caenis* | 1 | 0 | 0 | 0 | 0 | 0 | 0 | 0 | 0 | 0 | 0 | 0 | 0 | 0 | 0 | 0 |
| Ephemeroptera | Euthyplociidae | *Campylocia* | 7 | 8 | 0 | 18 | 9 | 1 | 7 | 1 | 29 | 15 | 17 | 53 | 44 | 56 | 12 | 17 |
| Ephemeroptera | Leptohyphidae | *Amanahyphes* | 0 | 0 | 0 | 0 | 0 | 0 | 3 | 1 | 5 | 5 | 2 | 2 | 0 | 5 | 0 | 5 |
| Ephemeroptera | Leptohyphidae | *Leptohyphes* | 7 | 0 | 0 | 0 | 0 | 0 | 0 | 0 | 0 | 0 | 0 | 0 | 0 | 0 | 0 | 0 |
| Ephemeroptera | Leptohyphidae | *Leptohyphodes* | 0 | 0 | 0 | 0 | 0 | 0 | 0 | 1 | 0 | 0 | 0 | 0 | 0 | 0 | 0 | 0 |
| Ephemeroptera | Leptohyphidae | *Traverhyphes* | 1 | 1 | 0 | 0 | 0 | 6 | 0 | 0 | 0 | 0 | 0 | 0 | 0 | 0 | 0 | 0 |
| Ephemeroptera | Leptohyphidae | *Tricorythodes* | 1 | 0 | 0 | 0 | 0 | 0 | 0 | 0 | 0 | 0 | 0 | 0 | 0 | 0 | 0 | 0 |
| Ephemeroptera | Leptohyphidae | *Tricorythopsis* | 0 | 0 | 0 | 0 | 0 | 0 | 0 | 0 | 0 | 0 | 1 | 0 | 0 | 4 | 0 | 0 |
| Ephemeroptera | Leptophlebiidae | *Askola* | 0 | 5 | 0 | 1 | 19 | 0 | 1 | 4 | 1 | 9 | 1 | 1 | 1 | 4 | 2 | 1 |
| Ephemeroptera | Leptophlebiidae | *Farrodes* | 112 | 16 | 6 | 45 | 22 | 84 | 25 | 22 | 29 | 29 | 26 | 4 | 14 | 19 | 5 | 124 |
| Ephemeroptera | Leptophlebiidae | *Hagenulopsis* | 6 | 4 | 6 | 0 | 1 | 1 | 8 | 73 | 0 | 22 | 0 | 0 | 2 | 8 | 0 | 55 |
| Ephemeroptera | Leptophlebiidae | *Hydrosmilodon* | 1 | 0 | 0 | 0 | 0 | 0 | 0 | 0 | 1 | 2 | 0 | 0 | 1 | 0 | 0 | 8 |
| Ephemeroptera | Leptophlebiidae | *Leentvaaria* | 1 | 0 | 0 | 0 | 0 | 0 | 0 | 0 | 0 | 0 | 0 | 0 | 0 | 0 | 0 | 0 |
| Ephemeroptera | Leptophlebiidae | *Miroculis* | 27 | 14 | 2 | 33 | 222 | 124 | 124 | 7 | 30 | 158 | 13 | 60 | 16 | 78 | 33 | 436 |
| Ephemeroptera | Leptophlebiidae | *Paramaka* | 13 | 0 | 0 | 0 | 0 | 2 | 1 | 1 | 0 | 0 | 0 | 0 | 0 | 0 | 0 | 0 |
| Ephemeroptera | Leptophlebiidae | *Simothraulopsis* | 0 | 0 | 0 | 0 | 0 | 0 | 0 | 0 | 3 | 5 | 0 | 3 | 1 | 0 | 0 | 4 |
| Ephemeroptera | Leptophlebiidae | *Terpides* | 0 | 2 | 6 | 2 | 14 | 0 | 3 | 0 | 0 | 0 | 0 | 0 | 0 | 0 | 0 | 0 |
| Ephemeroptera | Leptophlebiidae | *Ulmeritoides* | 2 | 1 | 0 | 5 | 49 | 15 | 28 | 4 | 0 | 0 | 0 | 0 | 0 | 0 | 0 | 0 |
| Ephemeroptera | Polymitarcyidae | *Campsurus* | 0 | 0 | 0 | 0 | 0 | 0 | 0 | 0 | 5 | 0 | 1 | 5 | 0 | 2 | 2 | 4 |
| Plecoptera | Perlidae | *Anacroneuria* | 172 | 201 | 22 | 26 | 66 | 63 | 65 | 32 | 4 | 21 | 62 | 22 | 42 | 48 | 17 | 30 |
| Plecoptera | Perlidae | *Macrogynoplax* | 2 | 2 | 4 | 6 | 4 | 32 | 30 | 17 | 18 | 14 | 74 | 24 | 59 | 19 | 20 | 23 |
| Trichoptera | Calamoceratidae | *Phylloicus* | 21 | 16 | 20 | 43 | 10 | 42 | 29 | 6 | 1 | 6 | 7 | 4 | 5 | 7 | 7 | 13 |
| Trichoptera | Ecnomidae | *Austrotinodes* | 0 | 0 | 0 | 0 | 0 | 0 | 0 | 0 | 2 | 1 | 0 | 2 | 0 | 10 | 1 | 0 |
| Trichoptera | Helicopsychidae | *Helicopsyche* | 1 | 0 | 0 | 0 | 0 | 0 | 0 | 0 | 0 | 7 | 3 | 11 | 7 | 31 | 3 | 13 |
| Trichoptera | Hydrobiosidae | *Atopsyche* | 3 | 1 | 0 | 0 | 0 | 0 | 0 | 0 | 0 | 0 | 0 | 0 | 0 | 0 | 0 | 0 |
| Trichoptera | Hydropsychidae | *Leptonema* | 6 | 25 | 4 | 2 | 3 | 16 | 7 | 16 | 3 | 1 | 66 | 31 | 44 | 91 | 9 | 19 |
| Trichoptera | Hydropsychidae | *Macronema* | 3 | 0 | 1 | 3 | 9 | 1 | 14 | 3 | 19 | 6 | 27 | 36 | 14 | 71 | 4 | 12 |
| Trichoptera | Hydropsychidae | *Macrostemum* | 0 | 0 | 5 | 0 | 0 | 0 | 0 | 1 | 5 | 17 | 32 | 7 | 0 | 15 | 1 | 0 |
| Trichoptera | Hydropsychidae | *Smicridea* | 10 | 33 | 3 | 0 | 0 | 1 | 0 | 0 | 2 | 0 | 23 | 6 | 15 | 0 | 1 | 2 |
| Trichoptera | Leptoceridae | *Amazonatolica* | 0 | 0 | 0 | 0 | 0 | 0 | 0 | 0 | 0 | 3 | 0 | 0 | 0 | 0 | 0 | 0 |
| Trichoptera | Leptoceridae | Genus A | 0 | 0 | 0 | 0 | 0 | 0 | 0 | 0 | 0 | 1 | 0 | 0 | 1 | 0 | 1 | 0 |
| Trichoptera | Leptoceridae | *Nectopsyche* | 27 | 849 | 2 | 0 | 0 | 28 | 0 | 1 | 0 | 2 | 4 | 1 | 2 | 8 | 2 | 0 |
| Trichoptera | Leptoceridae | *Notalina* | 1 | 0 | 0 | 0 | 0 | 0 | 0 | 0 | 0 | 0 | 0 | 0 | 0 | 0 | 0 | 0 |
| Trichoptera | Leptoceridae | *Oecetis* | 0 | 0 | 1 | 0 | 0 | 0 | 0 | 1 | 1 | 1 | 0 | 1 | 0 | 1 | 0 | 0 |
| Trichoptera | Leptoceridae | *Triplectides* | 2 | 1 | 0 | 1 | 6 | 3 | 3 | 0 | 1 | 1 | 0 | 0 | 0 | 1 | 1 | 3 |
| Trichoptera | Odontoceridae | *Marilia* | 0 | 0 | 0 | 0 | 2 | 0 | 0 | 0 | 2 | 9 | 1 | 6 | 4 | 0 | 6 | 1 |
| Trichoptera | Philopotamidae | *Chimarra* | 139 | 11 | 2 | 13 | 47 | 12 | 135 | 7 | 13 | 14 | 28 | 2 | 4 | 14 | 3 | 3 |
| Trichoptera | Polycentropodidae | *Cernotina* | 0 | 0 | 0 | 0 | 2 | 0 | 0 | 0 | 0 | 8 | 5 | 0 | 4 | 2 | 0 | 6 |
| Trichoptera | Polycentropodidae | *Cyrnellus* | 0 | 0 | 0 | 0 | 0 | 0 | 0 | 0 | 4 | 1 | 0 | 0 | 1 | 2 | 1 | 10 |
| Trichoptera | Polycentropodidae | *Polycentropus* | 0 | 0 | 0 | 1 | 0 | 0 | 1 | 0 | 0 | 0 | 0 | 0 | 0 | 0 | 0 | 0 |
| Trichoptera | Polycentropodidae | *Polyplectropus* | 0 | 3 | 2 | 0 | 1 | 0 | 0 | 0 | 0 | 3 | 6 | 3 | 1 | 2 | 0 | 0 |
| Trichoptera | Sericostomatidae | *Notidobiella* | 0 | 0 | 1 | 0 | 0 | 0 | 0 | 0 | 0 | 0 | 0 | 0 | 0 | 0 | 0 | 0 |
